# Particulate organic matter drives spatial variation in denitrification potential at the field scale

**DOI:** 10.1101/2023.11.20.567925

**Authors:** Emily R. Stuchiner, Wyatt A. Jernigan, Ziliang Zhang, William C. Eddy, Evan H. DeLucia, Wendy H. Yang

**Author notes:** Corresponding author: Emily R. Stuchiner. Open access: At this time, data are available upon request but will be uploaded to an appropriate repository pending manuscript acceptance.

## Abstract

High spatiotemporal variability in soil nitrous oxide (N_2_O) fluxes challenges quantification and prediction of emissions to evaluate the climate change mitigation outcomes of sustainable agricultural practices. Triggers for large, short-lived N_2_O emission pulses, such as rainfall and fertilization, alter soil oxygen (O_2_) and nitrate (NO_3_^−^) availability to favor N_2_O production via denitrification. However, the organic C (OC) needed to fuel denitrification may exhibit subfield variation that constrains the potential for high denitrification rates to occur, leading to spatial variation in N_2_O hot moments. We tested the hypothesis that the particulate organic matter (POM) fraction of soil organic matter controls subfield variation in denitrification potential by regulating availability of dissolved organic C (DOC), the form of OC used by denitrifiers. Among 20 soil samples collected across a maize field in central Illinois, USA, we found that potential denitrification rate was best predicted by POM C concentration (R^2^ = 0.35). Using multiple linear regression analysis that included other soil properties as explanatory variables, we found that POM C fraction of bulk soil (mg POM C g^−1^ SOC) was the most important predictor based on regression coefficient size (P < 0.01). Our results, which provide support for our hypothesis, suggest that consideration of the link between C and N cycling may be a key to predicting spatiotemporal variation in soil N_2_O emissions when denitrification is the dominant N_2_O source process.

## Introduction

The greenhouse gas nitrous oxide (N_2_O) is rapidly accumulating in the atmosphere due to human activities, such as nitrogen (N) fertilizer use to increase productivity in agroecosystems (Ravishankara et al., 2009, Cavigelli et al. 2012, Tian et al., 2020). High spatiotemporal variability in soil N_2_O emissions challenges quantification and prediction of emissions to evaluate the climate change mitigation benefits of sustainable agricultural practices (Groffman et al. 2009, Bernhart et al. 2015). Soil N_2_O production is highly sensitive to dynamic environmental drivers (Butterbach-Bahl et al. 2013, Wang et al. 2023), which can cause large, short-lived N_2_O emission pulses, referred to as hot moments, that disproportionately contribute to annual N_2_O budgets (Groffman et al. 2009, Anthony et al. 2023). At the same time, soil N_2_O emissions are often characterized by localized areas where above-average reaction rates occur, referred to as hot spots, even within homogenously managed agricultural fields (McDaniel et al. 2017, Krichels et al. 2019, Zhang et al. 2023 *in prep for Nature Geosciences*). Developing a predictive understanding of where N_2_O hot spots can occur when hot moments are triggered will improve our ability to account for spatiotemporal variability in soil N_2_O emissions in both measurement and modeling efforts.

Nitrous oxide hot moments occur when changes in the environment align to create conditions that stimulate high N_2_O production rates (Wagner-Riddle et al. 2020). Nitrogen fertilization and large rain events are considered the major triggers of nitrification and denitrification-driven hot moments in agricultural fields, respectively (Senbayram et al. 2012, Molodovskaya et al. 2012, Machado et al. 2020). Nitrification is typically only an important source of N_2_O following fertilization, whereas denitrification can be an important N_2_O source throughout the growing season (Ostrom et al. 2010, Harris et al. 2021, Stuchiner and von Fischer 2022a). Elevated soil NO_3_^−^ availability caused by nitrification or by direct fertilizer NO_3_^−^ inputs can support high rates of N_2_O production by denitrifiers, which reduce NO_3_^−^ through a series of enzymatic steps to N_2_O and N_2_ using organic carbon (OC) as the reductant (Firestone and Davidson 1989, Stuchiner and von Fischer 2022b). However, as an anaerobic process, denitrification contributes to N_2_O hot moments only when soil moisture conditions lead to the development of soil anoxia, such as following large rain events or spring thaw (Wagner-Riddle et al. 1998, Krichels et al. 2019). While fertilization and rainfall can uniformly supply N substrate and soil moisture to agricultural fields, they do not uniformly induce denitrification-driven N_2_O hot moments within fields (McDaniel et al. 2017, Krichels et al. 2019, Zhang et al. 2023 *in prep for Nature Geosciences*). This suggests that subfield variation in endogenous factors that regulate denitrification (e.g., O_2_, NO_3_^−^, OC) could constrain the potential for N_2_O hot moments to occur when exogenous events (e.g., fertilization, rainfall) alter environmental conditions to become favorable for N2O production.

Spatial variation in soil organic matter (SOM) may control the potential for denitrification-driven N_2_O hot moments at the subfield scale following large rain events and fertilization. Subfield variation in soil texture and topography can lead to spatial variation in soil drainage and NO_3_^−^ leaching (Basso et al. 2012, Zhu et al. 2015), which can contribute to subfield variation in N_2_O emissions (Turner et al. 2016, Lawrence et al. 2021). Large rain events should increase both soil moisture and microbial NO_3_^−^ accessibility enough to stimulate denitrification-driven N_2_O emissions across agricultural fields, unless the availability of SOM, the source of OC, constrains denitrification rates.

While the availability of SOM is ultimately modulated by where plant litter inputs accumulate in fields (Kravchenko et al. 2017), the accessibility of OC to microbes depends on whether the SOM resides in particulate organic matter (POM) or mineral-associated organic matter (MAOM). In general, POM exhibits faster turnover rates than MAOM, which is protected from decomposition by sorption to mineral surfaces or occlusion in soil micro-aggregates (Lavallee et al. 2020). Thus, the POM fraction of SOM could be more accessible to microbes than the MAOM fraction of SOM (Sokol et al. 2022), suggesting that it would be a more relevant OM pool to consider as a control on denitrification. In addition, regardless of pool size or OM quality, POM has been shown to serve as a more important source of dissolved OC (DOC) than MAOM to stimulate denitrification rates (Surey et al. 2021). Indeed, higher denitrification rates have been observed in regions of fields characterized by a greater abundance of crop residues, a precursor of POM (Li et al. 2016, Abalos et al. 2022a, Abalos et al. 2022b).

Here, we hypothesized that POM pool size controls subfield variation in denitrification potential by regulating DOC availability (Box 1). We predicted that soils containing more POM C would have higher denitrification potential because of greater OC accessibility to microbes. We measured potential denitrification rates in soil samples collected from 20 locations across a working farm in maize production in Champaign County, Illinois where we have previously observed spatial variation in N_2_O hot moments using autochamber measurements (Zhang et al. 2023 *in prep for Nature Geoscience*). We allowed native OC availability to constrain potential denitrification rates by omitting C amendments from the typical potential denitrification assay (Tiedje et al. 1994). We measured various indices of OC availability to microbes to relate to potential denitrification rates as explanatory variables, including POM C concentration, potential C mineralization rate, and DOC concentrations. We found that POM C was the most important explanatory variable predicting denitrification potential, providing support for our hypothesis.

**Box 1.**
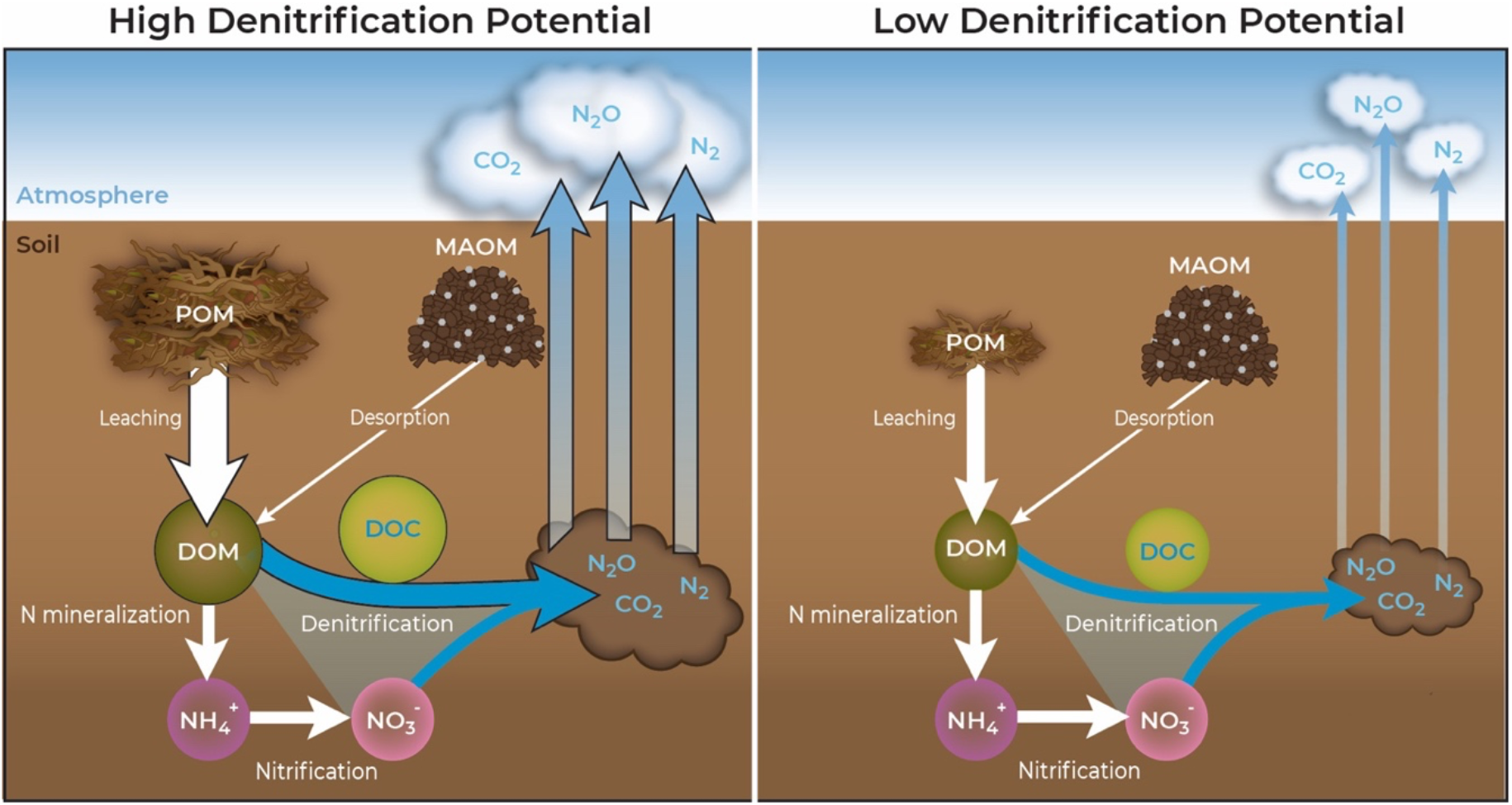
Conceptual diagram illustrating hypothesized role of the particulate organic matter (POM) pool size in regulating denitrification potential via leaching of dissolved organic matter (DOM) containing microbially accessible dissolved organic carbon (DOC). DOC is needed to fuel denitrification as the electron donor to reduce nitrate (NO_3_^−^) to nitrous oxide (N_2_O). Areas of fields with greater POM abundance therefore have high denitrification potential such that the onset of high soil moisture and NO_3_^−^ availability can trigger high N_2_O emissions (left panel). In contrast, areas of low POM abundance have low denitrification potential which constrain the response of N_2_O emissions to hot moment triggers (right panel). The relative sizes of icons and arrows represent approximate comparisons of pool sizes and process rates, respectively.

## Methods

### Study Site

Samples were collected from a working farm in Champaign County, IL (40.006°N, 88.290°W) that rotates annually between maize and soybean production. The farmer uses conservation tillage practices wherein vertical tillage occurs prior to planting in some but not all years. Urea ammonium nitrate (UAN) fertilizer is applied to the field during maize years shortly after planting at a rate of ∼200 kg N ha^−1^, and no fertilizer is applied during soybean years.

Over the period of 2014-2023 the mean annual air temperature at this site was 11 °C, with highest monthly mean temperature occurring in July (23.2 °C) and the lowest occurring in January (−3.5 °C, Midwestern Regional Climate Center 2023). The mean annual precipitation is 909 mm (Midwestern Regional Climate Center 2023). The soil at the farm consists of a roughly even combination of Drummer silty clay loam, Flanagan silt loam, and Catlin silt loam (Natural Resources Conservation Service).

### Experimental design

In this study, we took advantage of the spatial variation we observed in N_2_O fluxes across 20 autochamber locations at the farm during the 2021 growing season (Zhang et al. 2023 *in prep for Nature Geosciences*). Consistent N_2_O cold spots versus potential N_2_O hot spots suggested considerable subfield variation in denitrification potential, thereby providing an ideal setting for testing our hypothesis about controls on denitrification potential. In October 2022, we collected soils from the 20 locations, which are arranged into four distinct areas termed “nodes.” Each node consists of five autochambers spread across a ∼0.6 ha area within the overall study area, which spans ∼4.6 ha. Shortly after collecting the soil samples, we performed incubation assays and soil chemical measurements in the lab.

### Soil collection and storage

We collected soils in October 2022, approximately one week after harvest of the soybean crop. At each of the 20 autochamber locations we collected four soil cores to a depth of 30 cm using a 5-cm diameter soil auger. Two soil cores were bulked into each of two Ziploc bags, yielding two composite samples per autochamber location and 40 composite soil samples total.

Soil samples were kept on ice in the field to minimize microbial activity and were transported to the lab at the University of Illinois Urbana-Champaign (UIUC) within four hours for immediate homogenization and sieving. We sieved soils to 8 mm to retain some soil aggregate structure while removing large plant debris, roots, and rocks. Immediately after sieving, soil subsamples were extracted for chemical analyses and oven-dried at 105° C for determination of gravimetric water content (GWC). The soil samples were stored at 4° C for up to 12 hr before the denitrification enzyme assay (DEA; e.g., assay that measured potential denitrification rate) and C mineralization assay were initiated. The remaining soil samples were air-dried at room temperature (24° C) for several weeks prior to size fractionation, organic CN analysis, and soil texture analysis using the hydrometer approach as described by Gee and Bauder (1986).

### Denitrification enzyme assay

We used a modified denitrification enzyme assay (DEA) protocol to measure maximum denitrification rates under optimal NO_3_^−^ and O_2_ conditions for denitrification (i.e., excess NO_3_^−^ and anoxia) but with the constraint of native OC availability (i.e., no C amendment as usual). We weighed 25 g subsamples of sieved, fresh soil into 125 mL Wheaton bottles and brought the samples to room temperature over the course of ∼1 hr. To each Wheaton bottle, we added 25 mL of 1 mM KNO_3_^−^ solution in deionized (DI) water, capped the bottle, and then purged the headspace with N_2_ gas to induce anoxia. We then injected 99% acetylene gas (Airgas Industries, Illinois, USA) through a septum in the bottle cap into the bottle headspace to achieve 10% acetylene concentration by volume to inhibit N_2_O reduction. Each bottle was shaken vigorously by hand for 30 seconds and then gently shaken on an orbital shaker table for the duration of the 30-minute incubation period. We collected gas samples immediately after shaking (T0) and then every 10 minutes for 30 minutes, totaling four time points (T0, T10, T20, T30). Gas samples were analyzed for N_2_O concentration on a Shimadzu GC-2014 gas chromatograph equipped with an electron capture detector (Shimadzu Scientific, Illinois, USA). Since the change in N_2_O concentration over time for all samples was linear (R^2^ ≥ 0.997 in all cases), we calculated potential denitrification rates from the slopes of simple linear regression lines.

### C mineralization assay

We used a short-term C mineralization assay to estimate OC utilization by microbes as a proxy for OC availability to microbes. A pilot incubation of soil samples from our study site showed linear accumulation of CO_2_ in the jar headspace of soil samples incubated for one week, so we conducted the C mineralization assay over 72 hours. We weighed 30 g subsamples of sieved, fresh soil into 0.5 L Ball jars, loosely lidded the jars, and pre-incubated the soils at room temperature (24° C) for 24 hr. We pipetted DI water into the jars to bring the soil samples to 30% GWC and mixed the soil using a spatula to evenly distribute the added water. Following DI addition, the jars were sealed and immediately sampled to quantify the CO_2_ concentration at T0. Headspace gas sampling occurred again at 8, 24, 48, and 72 hr for CO_2_ flux determination over the 72-hour incubation period. Gas samples were stored for < 24 hr in pre-evacuated glass vials sealed with rubber septa and aluminum crimps. The gas samples were analyzed for CO_2_ concentration on the GC, which is also equipped with a thermal conductivity detector for CO_2_ analysis. Carbon mineralization rates were determined from the linear change in CO_2_ concentration during the incubation period.

### Soil chemical analyses

To support regression analyses to predict denitrification potential, DOC concentrations, and C mineralization rates, we measured a suite of soil chemical properties immediately after soil collection and immediately after the C mineralization assay (hereafter referred to as pre-incubation and post-incubation). We performed 2 M KCl extractions in a 5:1 ratio of KCl volume to dry soil equivalent mass for colorimetric analysis of NH_4_^+^ and NO_3_^−^ on a SmartChem 200 discrete analyzer (Unity Scientific, Brookfield CT, USA). We performed DI water extractions in a 3:1 ratio of DI volume to dry soil equivalent mass for DOC and TDN analysis on a Shimadzu Total Organic Carbon analyzer (Shimadzu TOC-L-CSH; Shimadzu Corp., Kyoto Japan) programmed to quantify non-purgeable OC and total N. Samples were vigorously shaken by hand for 30 seconds, centrifuged at 3000 rpm for 15 minutes, and then vacuum filtered through 0.7 µm filters that were ashed in a 450° C furnace. We performed 0.5 N HCl extractions in a 30:1 ratio of HCl volume to dry soil equivalent mass for colorimetric analysis of Fe(II) and Fe(III) using the ferrozine assay on a Genesys 20 spectrophotometer (Thermo Scientific Spectronic, MA). We also measured soil pH in slurries consisting of 5:1 ratio DI volume to fresh soil mass. Finally, we determined GWC for each pre-incubation sample by drying 5 g subsamples to a constant weight in a 105° C forced air oven. All extracts were frozen prior to analysis (within two weeks of extraction), except for the HCl extracts for Fe analysis which were stored at 4° C.

To test our hypothesis that spatial variation in POM leads to spatial variation in denitrification potential, we measured OC and TN concentrations in bulk soil as well as POM and MAOM size fractions. We present OC and TN in the POM and MAOM size fractions using two different measures: (1) as concentration relative to bulk soil mass, or (2) as bulk fraction relative to the bulk SOC or bulk soil TN (Table S1). For bulk soil, sieved soil samples were air-dried for four weeks and then oven-dried at 60° C for an additional 12 hr to remove residual moisture. Soils were ground to a fine powder using mortar and pestle, and then combusted for CN elemental analysis on a Elementar Vario Cube elemental analyzer (Hanau, Germany).

We separated POM and MAOM by size using the method described in Cotrufo et al. (2019). Briefly, the air-dried and 8 mm-sieved soils were sieved to 2 mm and then oven-dried at 60° C an additional 12 hr. The soil samples were then dispersed in 0.5% sodium hexametaphosphate and glass beads on a low-speed reciprocal shaker table for 18 hr. Dispersed soils were size-fractionated with a 53 µm sieve, and the > 53 µm fraction was washed with DI until the water ran clear. POM (> 53 µm) and MAOM (< 53 µm) fractions were dried at 60° C for four days and then weighed. Percent mass recovery ranged from 95% to 105% across all samples. The POM and MAOM samples were ground by hand using a mortar and pestle and then combusted for CN elemental analysis on a Velp 802 elemental analyzer (Velp Scientifica, New York, USA) at the Colorado State University EcoCore.

### Statistical analyses

Statistical analyses were performed in RStudio (version 4.2.2 (2022-10-31) -- “Innocent and Trusting” © 2022 The R Foundation for Statistical Computing). In all analyses, residuals were examined for departure from normality. All denitrification potential and C mineralization data were log-transformed to meet assumptions of normality in residuals, and all other data were normally distributed. During the DEA, one set of paired samples (i.e., the replicates from the same autochamber location) did not produce any N_2_O, presumably due to operator error while setting up the soil incubations. As such, these two samples were excluded from all statistical analysis, resulting in N = 38.

We calculated response ratios for soil properties that were measured before and after the C mineralization assay to understand the drivers of denitrification potential. Response ratios were quantified as the quotient of post-incubation over pre-incubation values. Thus, for ratios >1, the value increased during the C mineralization incubation, whereas for ratios <1, the value decreased during the C mineralization incubation. For the C mineralization assay, we held the soils in conditions that had previously demonstrated denitrification to be the dominant N_2_O-generating source process in the field (Zhang et al. 2023 *in prep for Nature Geosciences*). As such, measuring the response ratios of soil properties from the C mineralization assay allowed us to examine how differences in spatial drivers could trigger hot moments of denitrification under field-relevant conditions.

We used simple linear regressions to examine the drivers of the final DOC concentration (e.g., DOC post-incubation), and we used both simple and multiple linear regression to examine the drivers of denitrification potential. With simple linear regression we investigated the interaction between individual SOM pools and denitrification potential, and with multiple regression we predicted denitrification potential using all the measured soil variables as potential explanatory variables.

To avoid multicollinearity in the multiple linear regression model, prior to parameterizing the model, we performed principal components analysis (PCA) to identify which variables were co-correlated within three groups of data: soil/redox properties, N-related properties, and C-related properties. If two variables loaded along the same vector in the PCA, we removed the more weakly loaded variable from the multiple linear regression model (Figure S1). Through this procedure, we removed the NO_3_^−^-response ratio, DOC response ratio, and TDN response ratio because they correlated with %SON, MAOM C, and MAOM N, respectively. The MAOM C bulk fraction correlated with C mineralization rate, but both predictors were directly relevant to our research question, so we ran two multiple linear regression models, one parameterized without MAOM C bulk fraction and one parameterized without C mineralization rate. All remaining variables (not co-correlated), either response ratios or directly measured values, were included in the model, and we used backwards stepwise selection to identify the most parsimonious model. We also performed multiple linear regression analysis to predict C mineralization rate using the same approach as described for the denitrification potential models.

## Results

### Subfield variation in denitrification potential and soil properties

The 4.6 ha study area of the agricultural field exhibited considerable variation in denitrification potential and the soil properties that may control denitrification potential (Table 1). Denitrification potential spanned two orders of magnitude, from 3.95 to 338 ng N_2_O-N g^−1^ dry soil d^−1^. Soil NH_4_^+^ concentrations measured prior to the C mineralization incubation also spanned two orders of magnitude, from 0.05 to 8.8 µg N g^−1^ dry soil, whereas pre-incubation soil NO_3_^−^ concentrations varied only four-fold, from 1.7 to 6.8 µg N g^−1^ dry soil. Similarly, C mineralization rates varied six-fold, from 403 to 2396 mg CO_2_-C g^−1^ dry soil d^−1^, and bulk SOC, POM C, and MAOM C concentrations varied two-fold (Table 1). Bulk SOC ranged from 14.8 to mg C g^−1^ dry soil, with MAOM C consistently accounting for the majority of bulk SOC (89-96%). Water-extractable DOC measured prior to the C mineralization incubation also ranged three-fold from 18.8 to 47.8 µg C g^−1^ dry soil, with the ratio of post-to pre-incubation DOC varying from 0.58 to 1.

**Table 1.** Soil properties measured immediately after soil collection (pre-incubation) and at the end of the 72-hr C mineralization assay (post-incubation). In all cases, N = 38.

| Soil property | Pre-incubation |  |  | Post-incubation |  |  |
| --- | --- | --- | --- | --- | --- | --- |
| | Mean (± SE) | Minimum | Maximum | Mean (± SE) | Minimum | Maximum |
| Potential denitrification rate<br>($\text{ng N}_2\text{O-N g}^{-1}$ dry soil $\text{d}^{-1}$)* | | | | $39.2 \pm 9.52$ | 3.95 | 338 |
| C-mineralization rate<br>($\mu\text{g CO}_2\text{-C g}^{-1}$ dry soil $\text{d}^{-1}$) | | | | $733 \pm 60.5$ | 403 | 2400 |
| $\text{NH}_4^+$ ($\mu\text{g N g}^{-1}$ dry soil) | $1.48 \pm 0.341$ | 0.049 | 8.78 | $0.736 \pm 0.304$ | 0.019 | 9.69 |
| $\text{NO}_3^-$ ($\mu\text{g N g}^{-1}$ dry soil) | $4.58 \pm 0.238$ | 1.65 | 6.80 | $9.85 \pm 0.543$ | 5.34 | 22.9 |
| Fe (II) ($\mu\text{g-Fe g}^{-1}$ dry soil) | $10.1 \pm 0.621$ | 5.22 | 29.7 | $11.4 \pm 0.357$ | 7.98 | 17.0 |
| Fe (III) ($\mu\text{g-Fe g}^{-1}$ dry soil) | $471 \pm 13.7$ | 274 | 661 | $529 \pm 27.5$ | 248 | 823 |
| DOC ($\mu\text{g C g}^{-1}$ dry soil) | $27.0 \pm 0.961$ | 18.8 | 47.7 | $31.8 \pm 1.09$ | 19.6 | 43.9 |
| TDN ($\mu\text{g N g}^{-1}$ dry soil) | $14.8 \pm 0.640$ | 8.67 | 26.1 | $18.3 \pm 1.51$ | 7.94 | 52.1 |
| Soil pH | $6.42 \pm 0.062$ | 5.77 | 7.32 | $6.97 \pm 0.062$ | 6.29 | 7.74 |
| SOC ($\text{mg C g}^{-1}$ soil) | $22.2 \pm 0.676$ | 14.8 | 30.1 | | | |
| TN ($\text{mg N g}^{-1}$ soil) | $1.94 \pm 0.044$ | 1.48 | 2.39 | | | |
| POM C conc. ($\text{mg POM C g}^{-1}$ soil) | $1.60 \pm 0.057$ | 1.00 | 2.37 | | | |
| MAOM C conc. ($\text{mg MAOM C g}^{-1}$ soil) | $20.5 \pm 0.640$ | 13.3 | 28.2 | | | |
| POM N conc. ($\text{mg POM N g}^{-1}$ soil) | $0.130 \pm 0.007$ | 0.069 | 0.285 | | | |
| MAOM N conc. ($\text{mg MAOM N g}^{-1}$ soil) | $2.13 \pm 0.066$ | 1.34 | 3.07 | | | |
| Soil clay (%) | $21.9 \pm 0.342$ | 17.5 | 27.5 | | | |
| Soil sand (%) | $29.4 (\pm 1.64)$ | 8.22 | 54.0 | | | |

\*Potential denitrification rates were measured in a separate assay from the C mineralization assay.

### Predictors of subfield variation in denitrification potential

Simple linear regression suggested that POM C concentration best predicted subfield variation in denitrification potential. Particulate organic matter C concentration explained 35% of the variation in denitrification potential (p < 0.0001; Figure 1A). Denitrification potential also exhibited a positive but weaker relationship with water-extractable DOC measured after the 72-hour C mineralization assay (R^2^ = 0.19, p = 0.006; Figure 1B). In contrast, MAOM C concentration did not significantly correlate with denitrification potential (Figure 1C). Likewise, neither C mineralization rates nor pre-incubation DOC significantly correlated with denitrification potential (data not shown).

**Figure 1.**
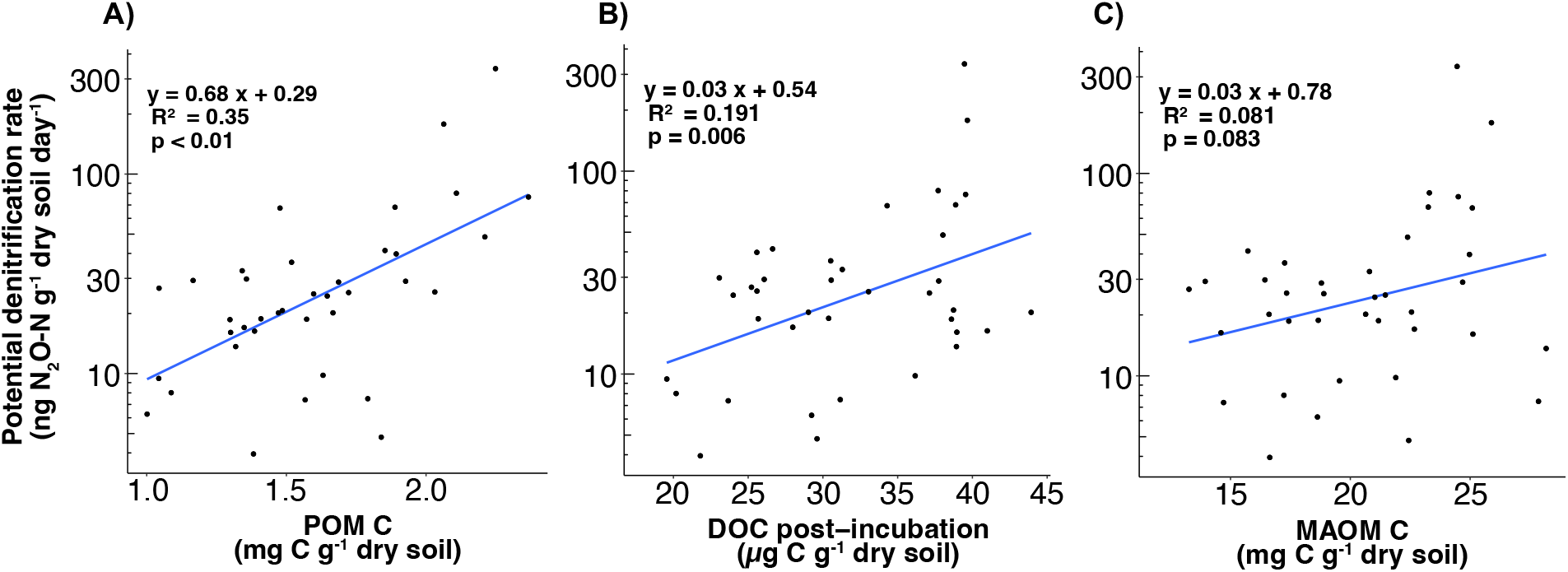
Simple linear regressions between log-transformed denitrification potential and concentrations of different OC pools, including (A) POM C, (B) water-extractable DOC at the end of the 72-hour incubation under moist soil conditions for the C mineralization assay, and (C) MAOM C. In all cases, N = 38.

Multiple linear regression analyses also showed that POM C was the best predictor of denitrification potential (Figure 2). In a model including MAOM C bulk fraction but not C mineralization rate, POM C bulk fraction had the largest regression coefficient (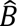 = 12.81) and was a highly significant predictor (p = 0.00182; Figure 2A). All other statistically significant explanatory variables in the model had small regression coefficients (ranging from −1.7 to 0.05), including MAOM C concentration, MAOM C bulk fraction, the ratio of NH_4_^+^ concentration from pre-incubation to post-incubation, and sand content (Figure 2B). In a model including C mineralization rate but not MAOM C bulk fraction, POM C bulk fraction still had the largest regression coefficient (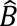 = 12.45) and was a highly statistically significant predictor of denitrification potential (p = 0.00304; Figure 2B). The % SOC and % sand were statistically significant, but again had small regression coefficients (0.67 and 0.01, respectively; Figure 2B).

**Figure 2.**
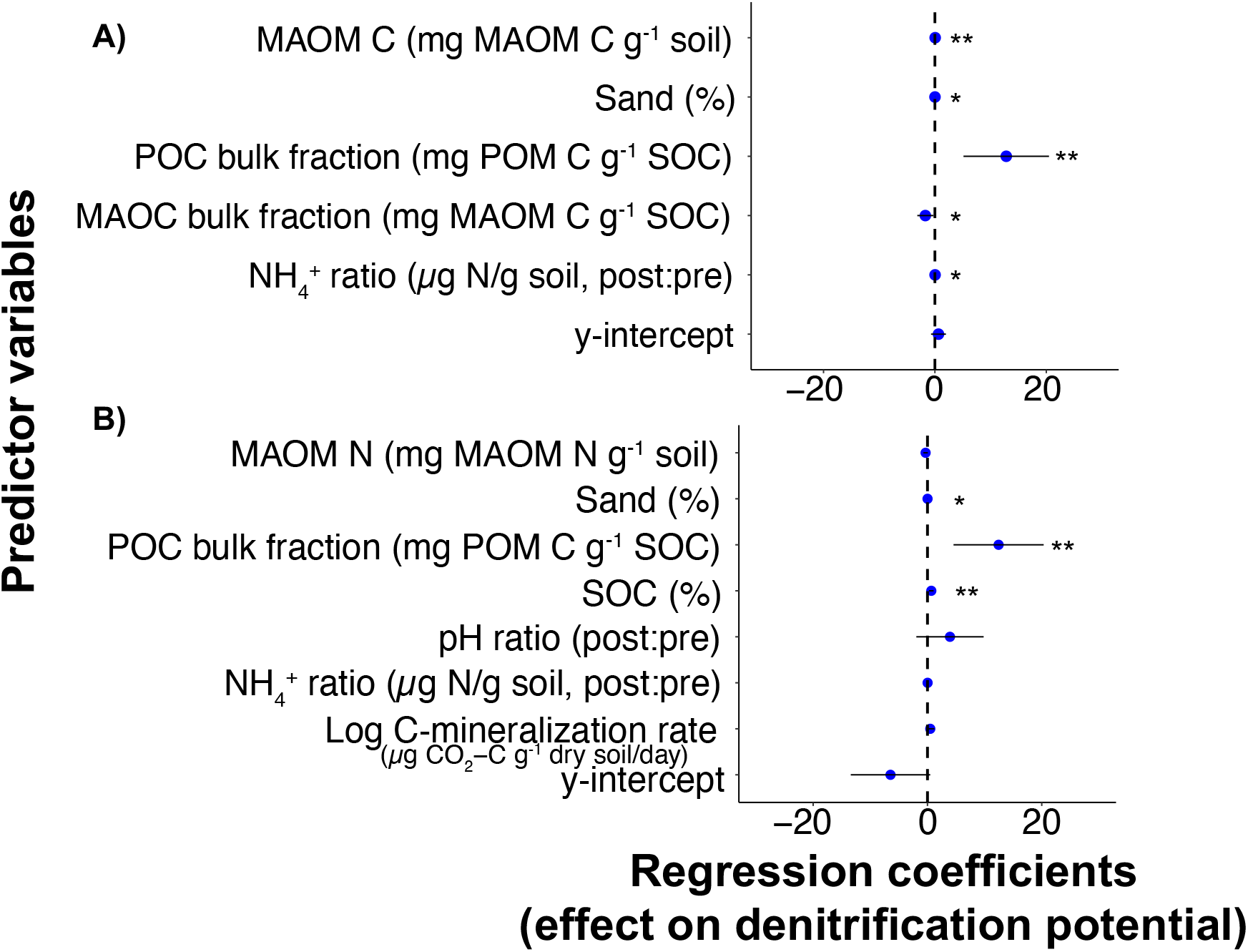
Plots of regression coefficients summarizing the multiple linear regression models of soil physical and chemical properties predicting denitrification potential. (A) shows the results of the multiple linear regression in which MAOM C bulk fraction was included in the model but C mineralization rate was not (R^2^ = 0.50), and (B) shows the results of the multiple linear regression in which C mineralization rate was included in the model but MAOM C bulk fraction was not (R^2^ = 0.57). In both models, N = 38, and bars represent ± 95% CI. Asterisks correspond to statistically significant correlations (** indicates p < 0.01, * indicates p < 0.05).

### Predictors of DOC concentrations

Pre- and post-incubation as well as the response ratio of water-extractable DOC concentrations were poorly predicted by all measures of POM C and MAOM C considered (only post-incubation data shown). Both POM C and MAOM C concentrations exhibited statistically significant but weak positive relationships with post-incubation DOC (R^2^ = 0.14 and R^2^ = 0.16, respectively, Table 2). None of the other POM C and MAOM C variables, including bulk fraction and C:N ratio, significantly correlated with post-incubation DOC concentrations (Table 2).

**Table 2.** Results from simple linear regressions for POM and MAOM properties as predictors of DOC concentration (mg C g^−1^ dry soil) after the 72-hour C mineralization assay incubation under moist conditions. Asterisks correspond to statistically significant correlations. In all regression models, N = 38.

| **Predictor of post-incubation DOC concentration** | **Equation** | **R<sup>2</sup>** | **p-value** |
| --- | --- | --- | --- |
| POM C concentration | $y = 7.14 x + 20.39$ | 0.14 | 0.02* |
| MAOM C concentration | $y = 0.69 x + 17.74$ | 0.16 | 0.01* |
| POM C bulk fraction | $y = 11.56 x + 30.96$ | 0.001 | 0.88 |
| MAOM C bulk fraction | $y = -2.55 x + 34.17$ | 0.001 | 0.86 |
| POM C:N | $y = -0.04 x + 32.29$ | < 0.001 | 0.92 |
| MAOM C:N | $y = 0.74 x + 24.65$ | 0.01 | 0.48 |

### Predictors of C mineralization rates

Carbon mineralization rates could not be predicted using any of the variables measured. After backwards stepwise selection, the multiple regression model included only two explanatory variables, POM C concentration and the post- to pre-incubation ratio of NH_4_^+^ concentrations (Figure 3); however, both variables had small effect sizes and were only marginally statistically significant (p = 0.06 and 0.08, respectively). In contrast, the y-intercept had a large positive effect size on C mineralization rate (p < 0.0001), indicating that most of the variation in C mineralization rate could not be explained by the variables we measured.

**Figure 3.**
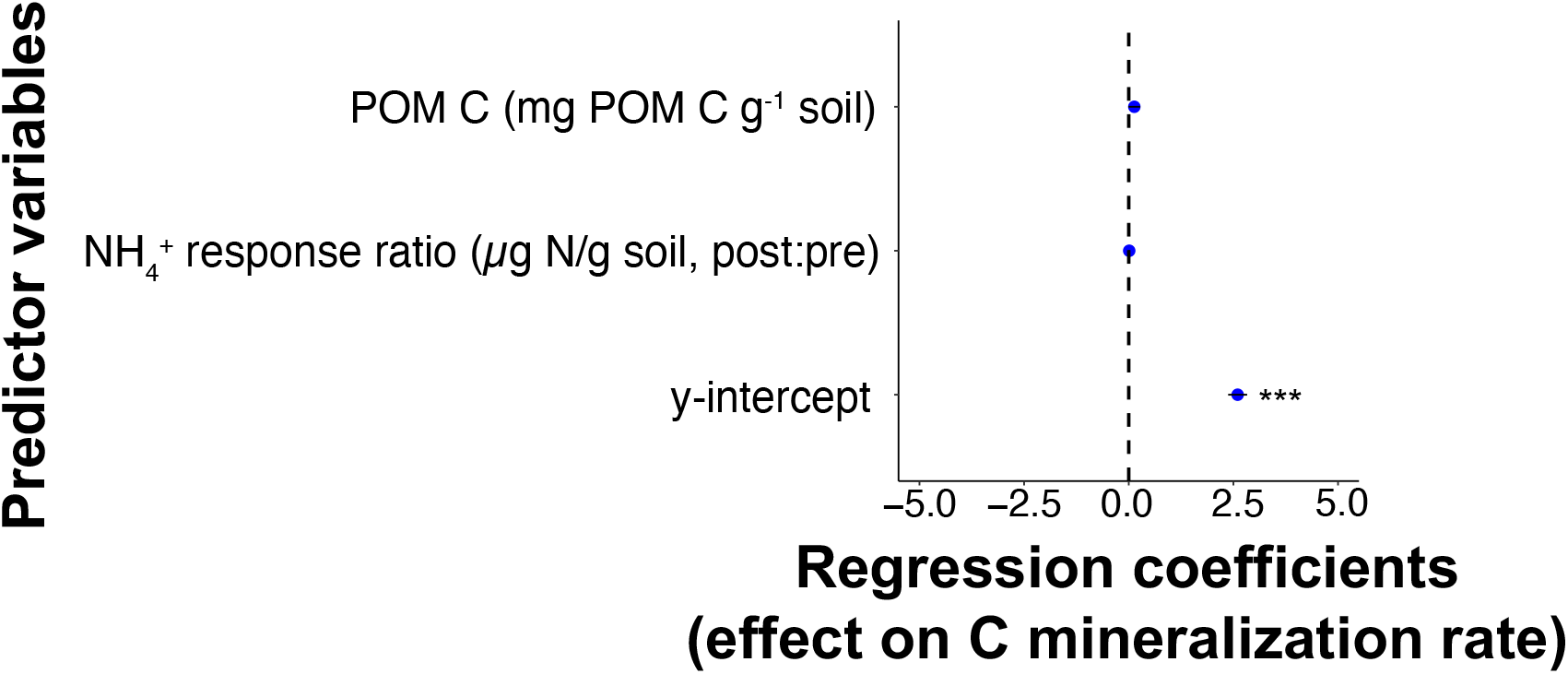
Effect size plot summarizing a multiple linear regression model of soil physical and chemical properties predicting C mineralization rates. In all cases, N = 38, and bars represent ± 95% CI. Asterisks correspond to statistically significant correlations (*** indicates p < 0.001).

## Discussion

Spatial and temporal variability in soil N_2_O emissions has persistently challenged measurement and modeling of N_2_O emissions to hinder N_2_O accounting and mitigation efforts (Barton et al. 2015, Lawrence et al. 2021). When changes in environmental conditions trigger high N_2_O emissions, N_2_O hot moments do not occur consistently even within homogenously managed agricultural fields (McDaniel et al. 2017, Krichels et al. 2019, Zhang et al. 2023 *in prep for Nature Geosciences*). By assaying soil samples from within a field that exhibited low versus high potential for N_2_O hot moments based on hourly N_2_O flux measurements (Zhang et al. 2023 *in prep for Nature Geosciences*), we found support for our hypothesis that spatial variation in denitrification potential at the subfield scale is determined by the accessibility of OC (Figure 1, Figure 2), a substrate that can limit denitrification rates when rainfall or fertilization depletes soil O_2_ or increases NO_3_^−^ supply, respectively (Box 1). Variables representing OC abundance in the POM pool of SOM were the strongest predictors of denitrification potential (Figure 1, Figure 2), which spanned over two orders of magnitude across the 20 sampling locations within 4.6 ha of a single agricultural field (Table 1). This suggests that OC from POM serves as a constraint on maximum denitrification rates under high soil moisture and NO_3_^−^. As such, understanding spatial variation in POM can help improve predictions of where high N_2_O emissions can occur within agricultural fields when hot moment triggers occur.

Particulate organic matter has faster C turnover than MAOM (Gentsch et al. 2015, Cotrufo et al. 2019, Lavallee et al. 2020), yet its functional importance in driving N cycling is underappreciated. While models are moving toward representing soil C cycling with separate POM and MAOM pools (Daly et al. 2021, Heckman et al. 2022), denitrification rates continue to be predicted based on bulk SOC properties (Yanai et al. 2003, Giltrap et al. 2010, Saha et al. 2021). Although MAOM C dominantly contributes to SOC across most sites (Sokol et al. 2022), including ours (Table 1), more OC can be leached from POM compared to MAOM (Surey et al. 2021). It is this DOC that is accessible for heterotrophic microbes such as denitrifiers to utilize because they must pass the OC across their cell membranes (Marschner and Kalbitz 2003). We did not find a correlation between DOC concentrations and denitrification potential (Figure 1B), and we did not see a correlation between DOC concentrations and microbial respiration either (Figure 3), likely because simultaneously high rates of DOC leaching and consumption confounded these relationships (Boddy et al. 2007, Gjettermann et al. 2008, Jones et al. 2009). Although we found that DOC concentrations were weakly correlated to both POM and MAOM (Table 2), the DOC derived from POM has been previously shown to stimulate denitrification more so than the DOC derived from MAOM (Surey et al. 2021). The strong positive correlations between denitrification potential and variables related to the POM pool support that POM represents the major source of OC accessible to denitrifiers (Figure 1, Figure 2). Furthermore, another high spatial resolution analysis conducted the prior year at our study site demonstrated a positive link between POM C and field-measured gross N_2_O production rates across the growing season (Zhang et al. 2023 *in prep for Nature Geosciences*). We therefore assert that predictions of spatial variation in soil N_2_O emissions can be improved by accounting for POM as the source of OC accessible to denitrifiers.

Understanding spatial variation in POM may be the key to improving predictions of where N_2_O hot moments occur, but this presents a new challenge. Whereas the spatial distribution of MAOM C is often positively correlated to soil clay content due to the role of clay minerals in adsorbing OC (Lavallee et al. 2020), POM C is more likely related to plant litter inputs (Kravchenko et al. 2017). In annual cropping systems where aboveground biomass is harvested, root biomass is the dominant plant litter input. However, given the difficulty in quantifying belowground productivity, root biomass data are sparse and characterized by high uncertainty due to small-scale spatial variation (Pausch and Kuzyakov 2018). Plasticity in root production in response to water availability can lead to variability in above- to belowground productivity that could manifest at the subfield scale with spatial variation in soil drainage (Gherardi and Sala 2020), so estimations of belowground productivity from aboveground productivity may be uncertain (Hui and Jackson 2006), making it difficult to predict POM spatial distribution from root biomass data. Nonetheless, in contrast to process-based predictions of spatial variation in POM C, machine learning algorithms trained on large datasets have been used to predict POM C at regional scales (Cotrufo et al. 2019, Lugato et al. 2021). Advances in streamlining SOM fractionation procedures to increase sample throughput to generate these types of datasets could help improve predictive modeling efforts of POM spatial distribution (Lugato et al. 2021).

Our findings have implications for understanding how agricultural management practices aimed at increasing SOC storage may have unintended effects on soil N_2_O emissions to counteract desired climate mitigation outcomes (Xia et al. 2018, Guenet et al. 2020). No-till or conservation-till cropping systems promote plant residues and subsequent SOM accumulation in croplands (Wang et al. 2020), leading to larger POM pools than in conventionally tilled systems (Six et al. 1999). Our study suggests that greater POM C can lead to greater soil N_2_O emissions. Indeed, denitrification rates have been shown to be higher in conservation tillage sites compared to conventional tillage sites (Mei et al. 2018, Ejack et al. 2020), although this depends on the crop residues present (Velthof et al. 2002). Although more labile residues can be readily metabolized by the soil microbial community and stabilized as MAOM (Cotrufo et al. 2013), many plant residues are primarily decomposed into POM before they are eventually turned over as microbial necromass to MAOM (Marucci et al. 2015, St. Luce et al. 2021). Accounting for the link between POM and denitrification will be necessary for improving predictions of climate mitigation outcomes from agricultural management practices.

In conclusion, we have demonstrated that OC accessibility via POM can help explain spatial and temporal variability in soil N_2_O emissions in an agricultural field that exhibits consistent N_2_O cold spots and potential N_2_O hot spots (Zhang et al. 2023 *in prep for Nature Geosciences*). Of the three major factors controlling denitrification rates (i.e., soil moisture, NO_3_^−^, and OC), only OC is constrained to endogenous sources within the ecosystem. Therefore, when exogenous inputs of rainfall or fertilizer increase soil moisture or NO_3_^−^ to stimulate denitrification rates, the endogenous supply of OC largely from POM can limit denitrification rates to create spatial variation in soil N_2_O emissions. This connection between C and N cycling may be a key to predicting spatial and temporal variation in soil N_2_O emissions when denitrification is the dominant N_2_O source process.

## Acknowledgements

We would like to thank Michelle Haddix and the Cotrufo lab for their generous assistance with soil size fractionations. We would also like to thank the EcoCore staff at Colorado State University for allowing us to use their Elemental Analyzer. None of this work would have been possible without the generous field and laboratory support from the Yang lab research specialists Chloe Yates and Haley Ware. This work was funded with support from the DOE ARPA-E Smartfarm Program.

