## Supplement for "Particulate organic matter drives spatial variation in denitrification potential at the field scale"

### Contents of this file

Table S1
Figure S1

**Table S1.** Measures of organic carbon (OC) and total nitrogen (TN) in the particulate organic matter (POM) and mineral-associated organic matter (MAOM) size fractions relative to bulk soil. Each variable will be referenced in the text by its name in the "Variable" column. The Equation column specifies how the value for each variable was calculated. % POM C and % MAOM C are OC concentrations relative to POM and MAOM mass, respectively, as determined by elemental analysis of the size fractions. Similarly, % POM N and % MAOM C are TN concentrations relative to POM and MAOM mass, respectively, as determined by elemental analysis of the size fractions.

| Variable | Units | Meaning | Equation |
|---|---|---|---|
| POM C concentration | mg POM C/g soil | POM C concentration in bulk soil | (1) mg POM C $g^{-1}$ soil = ((% POM ÷100) * (% POM C ÷ 100)) * 1000 |
| POM C bulk fraction | mg POM C/g SOC | Fraction of SOC that is in POM C | (1) mg C $g^{-1}$ soil = (% SOC ÷ 100) * 1000<br>(2) POM C bulk fraction = mg POM C $g^{-1}$ soil ÷ mg C $g^{-1}$ soil |
| POM N concentration | mg POM N/g soil | POM N concentration in bulk soil | (1) mg POM N $g^{-1}$ soil = ((% POM ÷100) * (% POM N ÷ 100)) * 1000 |
| POM N bulk fraction | mg POM N/g TN | Fraction of TN that is in POM N | (1) mg N $g^{-1}$ soil = (% SON ÷ 100) * 1000<br>(2) POM N bulk fraction = mg POM N $g^{-1}$ soil ÷ g N $g^{-1}$ soil |
| MAOM C concentration | mg MAOM C/g soil | MAOM C concentration in bulk soil | (1) mg MAOM C $g^{-1}$ soil = ((% MAOM ÷100) * (% MAOM C ÷ 100)) * 1000 |
| MAOM C bulk fraction | mg MAOM C/g SOC | Fraction of SOC that is in MAOM C | (1) mg C $g^{-1}$ soil = (% SOC ÷ 100) * 1000<br>(2) MAOM C bulk fraction = mg MAOM C $g^{-1}$ soil ÷ g C $g^{-1}$ soil |
| MAOM N concentration | mg MAOM N/g soil | MAOM N concentration in bulk soil | (1) mg MAOM N $g^{-1}$ soil = ((% MAOM ÷100) * (% MAOM N ÷ 100)) * 1000 |
| MAOM N bulk fraction | mg MAOM N/g TN | Fraction of TN that is in MAOM N | (1) mg N $g^{-1}$ soil = (% SON ÷ 100) * 1000<br>(2) MAOM N bulk fraction = mg MAOM N $g^{-1}$ soil ÷ g N $g^{-1}$ soil |

**Figure S1.** Principal component analysis (PCA) of the correlations among predictor variables of potential denitrification rate. Variables were grouped by soil/redox properties (Panel **A**), C properties (Panel **B**), and N properties (Panel **C**). If two variables loaded along the same vector in the PCA, we removed the weaklier loaded variable from the multiple linear regression model used in the main text analysis that examined how the predictors affected denitrification potential.

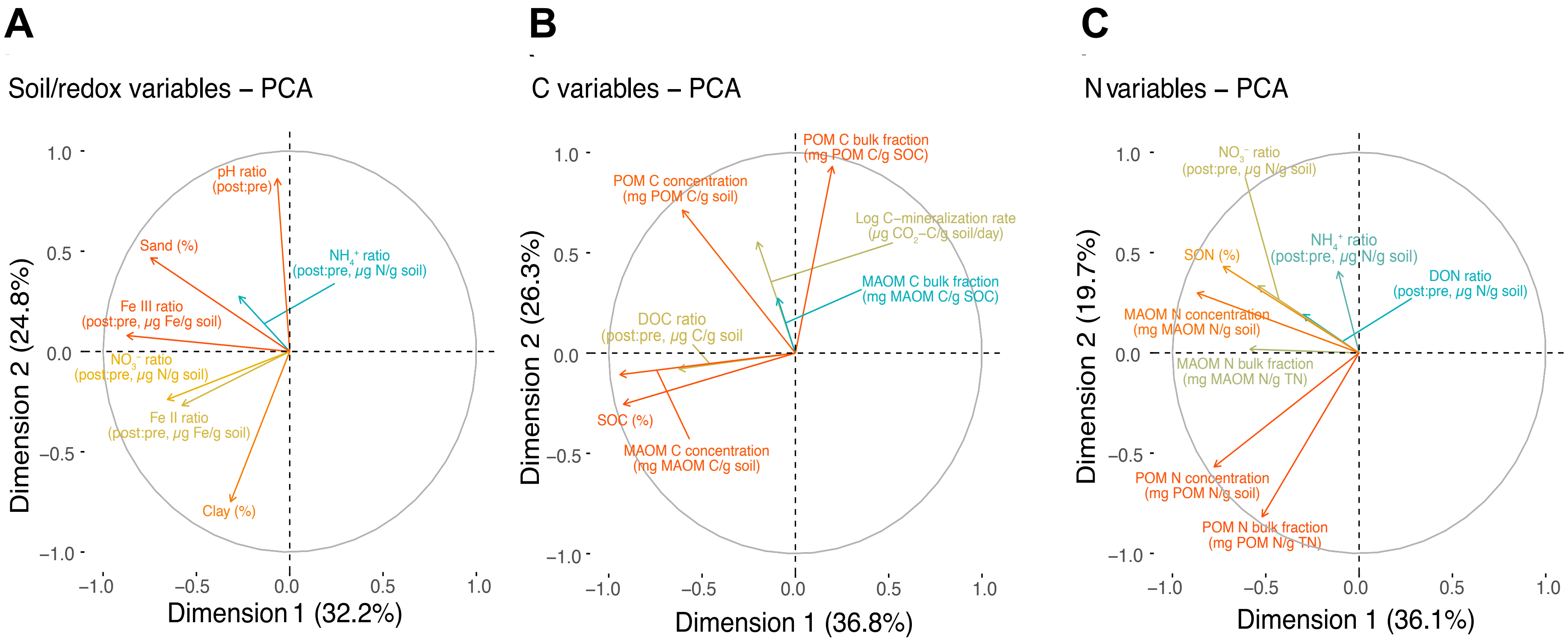
